# Identification and Integration of Key-Metabolic Reactions from Untargeted Metabolomics

**DOI:** 10.1101/2023.05.15.540613

**Authors:** Nikolai Köhler, Vivian Würf, Tim D. Rose, Josch K. Pauling

## Abstract

Metabolomics has become increasingly popular in biological and biomedical research, especially for multi-omics studies, due to the many associations of metabolism with diseases. This development is driven by improvements in metabolite identification and generating large amounts of data, increasing the need for computational solutions for data interpretation. In particular, only few computational approaches directly generating mechanistic hypotheses exist, making the biochemical interpretation of metabolomics data difficult. We present *mantra*, an approach to estimate how metabolic reactions change their activity between biological conditions without requiring absolute quantification of metabolites. Starting with a data-specific metabolic network we utilize linear models between substrates and products of a metabolic reaction to approximate deviations in activity. The obtained estimates can subsequently be used for network enrichment and integration with other omics data. By applying *mantra* to untargeted metabolomics measurements of Triple-Negative Breast Cancer biopsies, we show that it can accurately pinpoint biomarkers. On a dataset of stool metabolomics from Inflammatory Bowel Disease patients, we demonstrate that predictions on our proposed reaction metric generalize to an independent validation cohort and that it can be used for multi-omics network integration. By allowing mechanistic interpretation we facilitate knowledge extraction from metabolomics experiments.

## 1 Introduction

Metabolites display the product of metabolism and thereby the metabolic state of an organism. Their chemical structures are as diverse as their functions, ranging from pure energy metabolism to immune modulation and environmental sensing [1]. Owing to the essential nature of many of these functions, metabolism is tightly regulated through the control of enzymatic activity. This regulation can happen on different levels, such as the amount of enzyme, post-translational modifications, or allosteric regulation by other metabolites [2, 3, 4, 5, 6]. Since metabolite concentrations are also highly influenced by environmental factors such as diet, medication, or a host’s microbiome [7], the metabolic phenotype is the highly complex result of internal metabolic processes and environmental factors. The resulting metabolic phenotype is often referred to as the “metabotype” [8].

Metabolomics, the large-scale study of metabolites, is used to characterize the changes in metabolite levels. Due to the importance of metabolic processes for almost any aspect of life, metabolomics is becoming increasingly popular for biological and biomedical research. Even though the chemical analysis of metabolites, most commonly via Mass Spectrometry (MS), has made great progress in the last decade confident large-scale identification remains difficult, and especially reliable absolute quantification is only possible for a small set of target metabolites. These shortcomings make it particularly challenging to apply metabolomics for exploratory purposes with no clear hypothesis when trying to understand the molecular mechanisms behind different biological conditions.

In addition to such analytical challenges, computational metabolomics, outside molecule identification, is still a small field. Consequently, a rather small number of methods for the computational interpretation of metabolomics data are available. The most commonly used analyses are “classical” univariate statistical tests and fold-changes as well as multivariate approaches such as Principal Component Analysis (PCA) and Partial Least Squares(-Discriminant Analysis) (PLS(-DA)) [9]. While these methods allow it to extract significantly altered metabolites and get an overview of how different the metabolome in different conditions is, they do not allow for direct biochemical interpretation of the results. Instead, specific over-representation or pathway enrichment methods, e.g. MSEA [10] or IMPaLA [11], are used to obtain high-level summaries. Despite delivering biochemically more coarse-grained and comprehensible results, they don’t allow for the generation of mechanistic hypotheses on the level of *de-novo* pathways or quantitatively for individual reactions.

To computationally propose such mechanistic interpretation, metabolic networks can be utilized. They can be represented as directed bipartite graphs in which metabolites and reactions are nodes connected by (directed) edges - substrate or product relations - which are catalyzed by specific enzymes [12]. Such networks are available for many organisms nowadays, e.g. from KEGG [13] or BioCyc [14], but only cover parts of the entire metabolome, therefore limiting the scope to known metabolic reactions.

One way to leverage such networks is metabolic modeling, more precisely kinetic or constraint-based modeling [15]. The advantage of such methods is that they are able to make precise predictions on how metabolism behaves, given that the underlying model assumptions are valid. However, this dependence is also a major weak point since, particularly for eukaryotes, the correct parameterization of these models is hard, yet critical for the predicted outcome to the extent of yielding possibly contrasting results [16].

Another strategy to incorporate prior-knowledge networks into data analysis is to use graph theoretic approaches. Especially network enrichment, which aims at identifying subgraphs characterized by high changes between conditions, has been extensively used and studied in genomics, transcriptomics, and proteomics [17]. For metabolomics, only a few such methods are available [12]. One of them is the MetaboRank algorithm [18]. It turns the metabolic network into a Markov model by defining transition probabilities between substrate-product pairs on the basis of the proportion of mapped atoms. The “metabolic fingerprint”, as the authors call it, is then computed by performing random walks via a variation of Personalized Page Rank [19]. Another method also using random walks/diffusion to assign relevance to entities in a metabolic graph was introduced by Picart-Armada et al. [20]. In contrast to using metabolic networks, this approach uses a KEGG graph including different hierarchies from compounds down to the level of modules and pathways. This makes it one of the few methods that gives results on metabolic reactions directly. To compute the relevance of each node in the graph, heat diffusion is simulated with only significantly altered metabolites being able to introduce heat into the system. Each node’s relevance is then determined by the heat flow going through or the proportion of random walks including it and then compared to the outcome of a null model in which significances are randomly permuted to avoid structure-based biases.

One drawback of the available methods is, that they either only consider the network-topological properties of metabolites or incorporate only significances, which solely rely on univariate tests and a p-value threshold. In this work, we tackle these shortcomings by presenting an approach we named *mantra* (**M**et**a**bolic **N**e**t**work **R**eaction **A**nalysis), to estimate how the activity of individual metabolic reactions changes between biological conditions on the basis of relative metabolite intensities. In contrast to existing graph-based metabolomics methods, our approach does not rely on computing univariate statistics for all metabolites but uses metabolite abundances to obtain samplewise estimates. It thereby avoids the need for a significance threshold prior to enrichment and allows for the integration of additional omics layers via their biochemical connections.

Using two independent clinical studies, we demonstrate how our approach conserves a significant proportion of the original variation while enabling the generation of hypotheses on the metabolic mode of action. Furthermore, we show the capability of our method to improve the integration of metabolomics into a multi-omics context and how it can be used to define *de-novo* disease pathways. These characteristics make it a promising concept for advancing the mechanistic interpretation of metabolomics data in a clinical context. The idea presented can be used as a basis to develop a new class of approaches for metabolic network analysis.

## 2 Results

We first start by briefly giving an overview of our approach before presenting its application on independent clinical data sets. A more detailed explanation of each step of the workflow is given in Materials & Methods.

The input to the proposed method is simply a table of normalized metabolite intensities and, optionally, a metabolic network. If no metabolic network is given, the metabolites are first mapped onto internal identifiers to generate a network from a custom database built from KEGG [13], Reactome [21] and Virtual Metabolic Human (VMH) [22]. Its structure is schematically shown in the top right of Figure 1. To estimate the activity of each reaction in the network, a linear model with the substrates as the predicting and the products as the dependent variables is computed. Next, the explained variance for each sample is computed from the distance of the predicted product values to the actual product values. The intuition behind this step is that the higher the activity of a metabolic reaction, the higher the dependence structure between substrate and product concentrations (and thus the measured intensities) will be. If a reaction now changes its activity, the association persists, but the coefficients describing it will likely change. Therefore, this alteration can be observed in a change of residuals. These values can now directly be used to perform network enrichment where the objective is to find a connected subgraph in which the difference between residual distribution is maximized for two given sample groups. Since our proposed metric is per-sample, it can also be used to correlate them to features from other omics layers, for example, microbial abundances. The associations can also be restricted to known interactions if provided with the metabolic network or provided at network generation when using our custom database. To avoid assumptions on distributions these interactions are computed using Spearman’s rank correlation.

**Figure 1:**
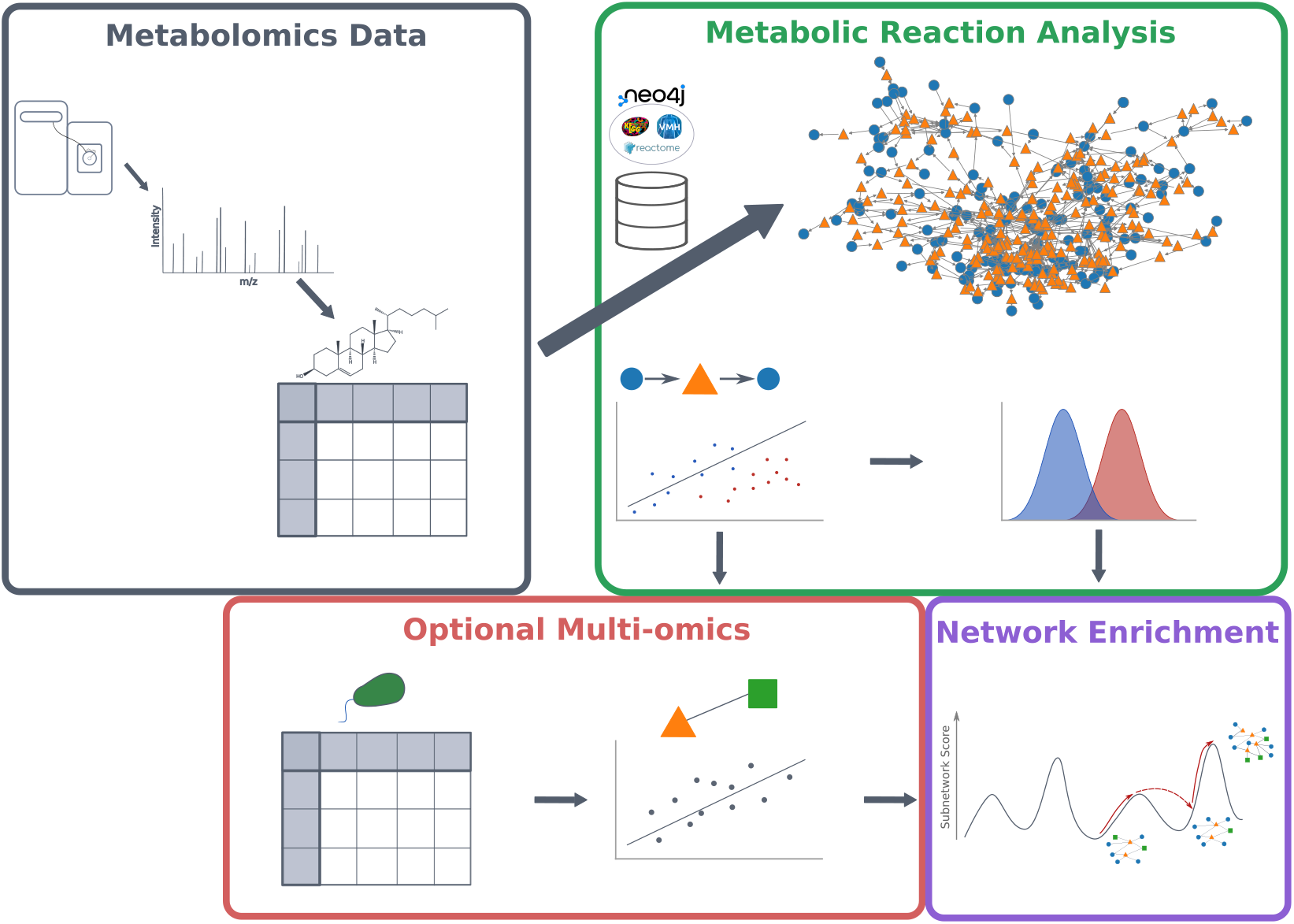
Overview of the *mantra* workflow. Starting with identified (untargeted) metabolomics data (*gray box*), a metabolic network containing only measured metabolites is constructed. It is a directed bipartite graph with metabolites depicted as blue circles and metabolic reactions as orange triangles. For each metabolic reaction in this network, the reaction activity relative to the activity in the control group is estimated using a linear model between the substrate and product abundances (*green box*). These activities can either be directly used for network enrichment (*purple box*) or together with multi-omics data (*red box*). In the latter case, the estimated per-sample activity values are correlated with the expression of each feature in the multi-omics data.

### 2.1 *mantra* Recovers Known Key Reactions in Triple-Negative Breast Cancer

To demonstrate the capabilities of our approach to metabolomics, we chose a data set by Xiao et al. [23]. The processed data contains 330 Triple-Negative Breast Cancer (TNBC) and 149 control samples, and 594 identified metabolites with their corresponding intensities. Mapping the metabolites onto our database yielded a network with 173 metabolites and 254 reactions (Figure 2a). The reduced number of metabolites is due to metabolites not being matched to database identifiers present in our database or not being connected to any reaction for which at least one substrate and one product were measured.

**Figure 2:**
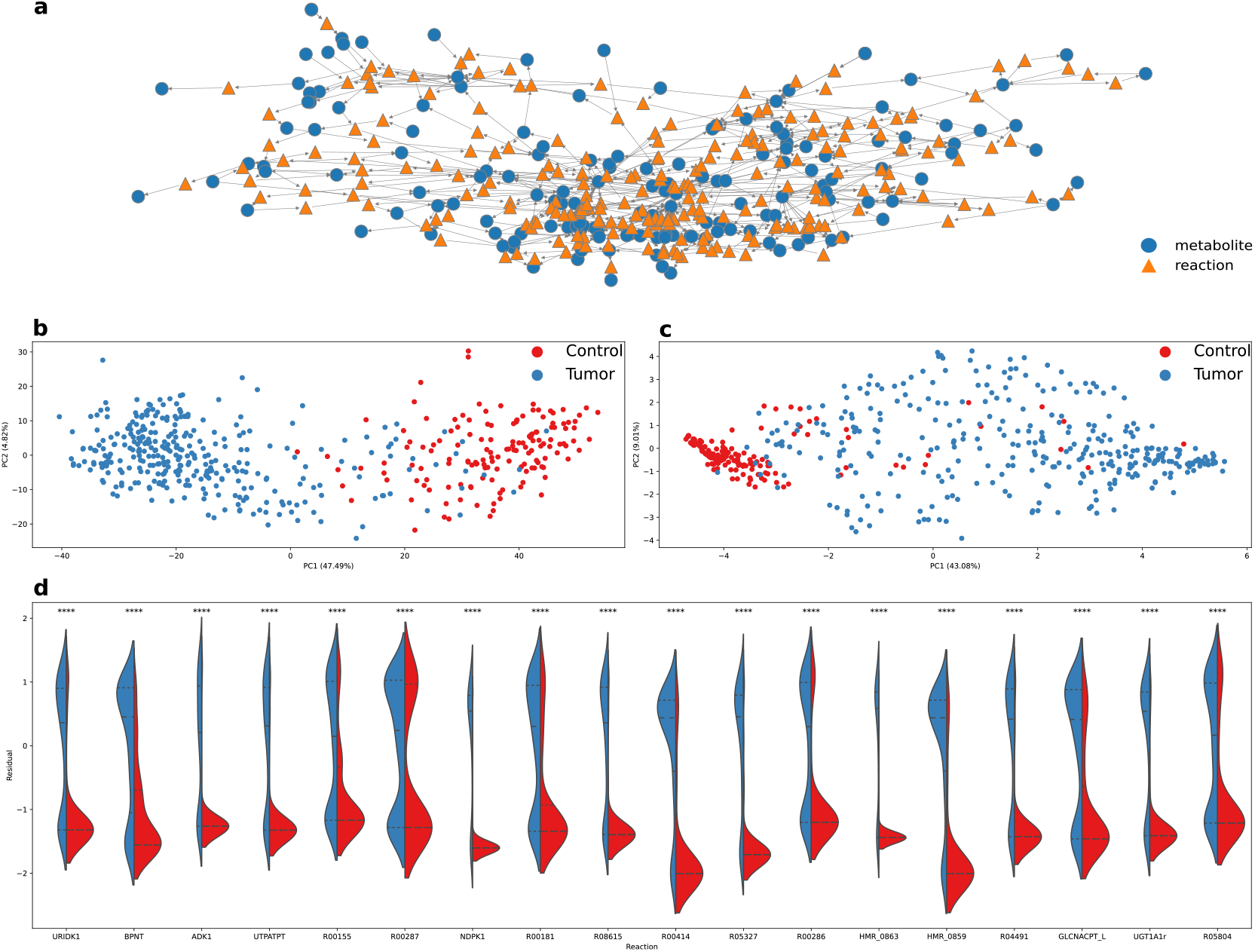
*mantra* results on Control vs. Triple-Negative Breast Cancer (TNBC). **a** Metabolic network build with the metabolites measured and identified in Xiao et al. [23]. Metabolites are shown as blue circular nodes, reactions as orange triangles. **b** PCA of the processed metabolite data with samples colored by condition. Despite a few overlapping samples, the groups are clearly separated by PC1, which explains almost half of the variance in the data. **c** PCA of the reaction activity estimates calculated by *mantra* with samples colored by condition. Similar to *b*, sample groups are separated by PC1, with only a minor reduction in the fraction of explained variance. In contrast to the original metabolome data, control samples show a reduced within-group variance, while tumor samples show a higher within-group variance. **d** Distributions of activity estimates for the most significantly changing reactions. Significance values were computed with a Wilcoxon rank sum test and Bonferroni correction. ^****^ indicates a corrected p-value < 0.001.

Most nodes in the network have between 2 and 5 connections (Supplementary Figure S1). Metabolic reaction nodes do not range higher than this, which is expected, considering that reactions generally don’t have a large number of substrates and products. For metabolite nodes, however, there are some hub nodes, which take part in up to 50 metabolic reactions. Metabolites with a node degree above 10 are listed in Supplementary Table S1.

To evaluate how well our proposed metric for relative reaction activity conserves the variance contained in the original metabolomics data, their PCA plots are shown in Figure 2b (metabolome data) and c (reaction data). Sub-figure b shows a separation of the control and the TNBC samples along PC1, which explains around 47% of the total variance. In comparison, PC1 of the reaction data in Figure 2c explains a little less variance. Although the variance between control samples is low, there are few outliers on the far right. The TNBC group, on the other hand, shows a much higher variation than in the general metabolome data. Nevertheless, Figure 2c shows that our approach is able to retain biological variance and separate the clinical conditions.

Going into a more detailed analysis on which metabolic reactions are identified as the most changing between conditions, 18 reaction nodes appear as highly significant (Figure 2d). Some of these reaction nodes represent more than one metabolic reaction, as some reactions have the same measured substrates and products, making the possibly involved catalytic enzymes indistinguishable. The details of the reactions from Figure 2d are given in Supplementary Table S2.

Most of these reactions involve nucleotides, most prominently Uridine Mono-Phosphate (UMP) and Uridine Di-Phosphate (UDP), and different glycosylation reactions. Both of these reaction classes, but especially UDP-related reactions have been identified as key players in breast cancer [24, 25, 26, 27, 28, 29]. Notably, enzymes catalyzing 4 of the significant reactions - UDP-glucuronosyltransferase (UGT), UMP-CMP kinase (CMPK1) and UDP-glucose-dehydrogenase (UGDH) - are known to be associated with breast cancer risk or are significant prognostic biomarkers [24, 19, 26, 27]. Furthermore, Adenylate Kinase 4 was found to regulate resistance to Tamoxifen treatment [29], which is used to treat Estrogen Receptor (ER) positive breast cancer patients. This is especially interesting since UGT enzymes conjugate ER ligands, forming a direct link between these reactions on a signaling level. These findings indicate that our approach is able to generate both usable and testable hypotheses on changes in metabolic activity.

In addition to this non-metabolic link between significant reactions, the reactions from Figure 2d form a connected subgraph of the metabolic network in Figure 2a, shown in Supplementary Figure S2. This supports our hypothesis, that in addition to metabolic networks being small-world networks, changes in metabolic activity are often constrained to a smaller subgraph of metabolic reactions, despite changes on metabolite level usually being observed throughout a larger part of the graph. Linking dysregulation of individual reactions back to broader metabolic mechanisms additionally allows one to elucidate mechanistic relations on a higher level. Because the computed activity values are computed for each sample, it is also possible to identify sub-populations within the group of disease samples. In combination with the mechanistic character of our method, this enables insights into metabolic alterations down to a patient-specific level making it a promising tool for precision medicine applications.

### 2.2 Application to Inflammatory Bowel Disease and Multi-omics Integration

The second analysis we present here is applying the *mantra* approach to data from the PRISM cohort from Franzosa et al. [30]. It consists of 155 stool samples, 34 control and 121 Inflammatory Bowel Disease (IBD) samples, which are grouped into 68 Crohn’s Disease (CD) and 53 Ulcerative Colitits (UC) samples. For the presented analysis only control and CD samples were used. After filtering and mapping the metabolites and microbial species onto the internal database (for details see Materials & Methods), 138 metabolites and 70 microbial species were retained and 108 metabolic reactions were included in the metabolic network.

#### Linking Metabolomics-based Reaction Activity to Differential Enzymatic Potential

An overview of the variance in the processed metabolome data and the computed reaction activity data are shown using PCA in Figure 3 c and d, respectively. Generally, both spaces seem to discriminate the patient groups well. Sub-figure c shows a good separation of CD and control samples along PC1, which explains around 20% of the total variance. Notably, CD samples appear to be metabolically more diverse than control samples. While the separation between CD and control along PC1 in based sub-figure d, which is based on our proposed reaction activity estimates, is a little less clear with some control outliers, the explained variance along this component is increased to around 33%.

**Figure 3:**
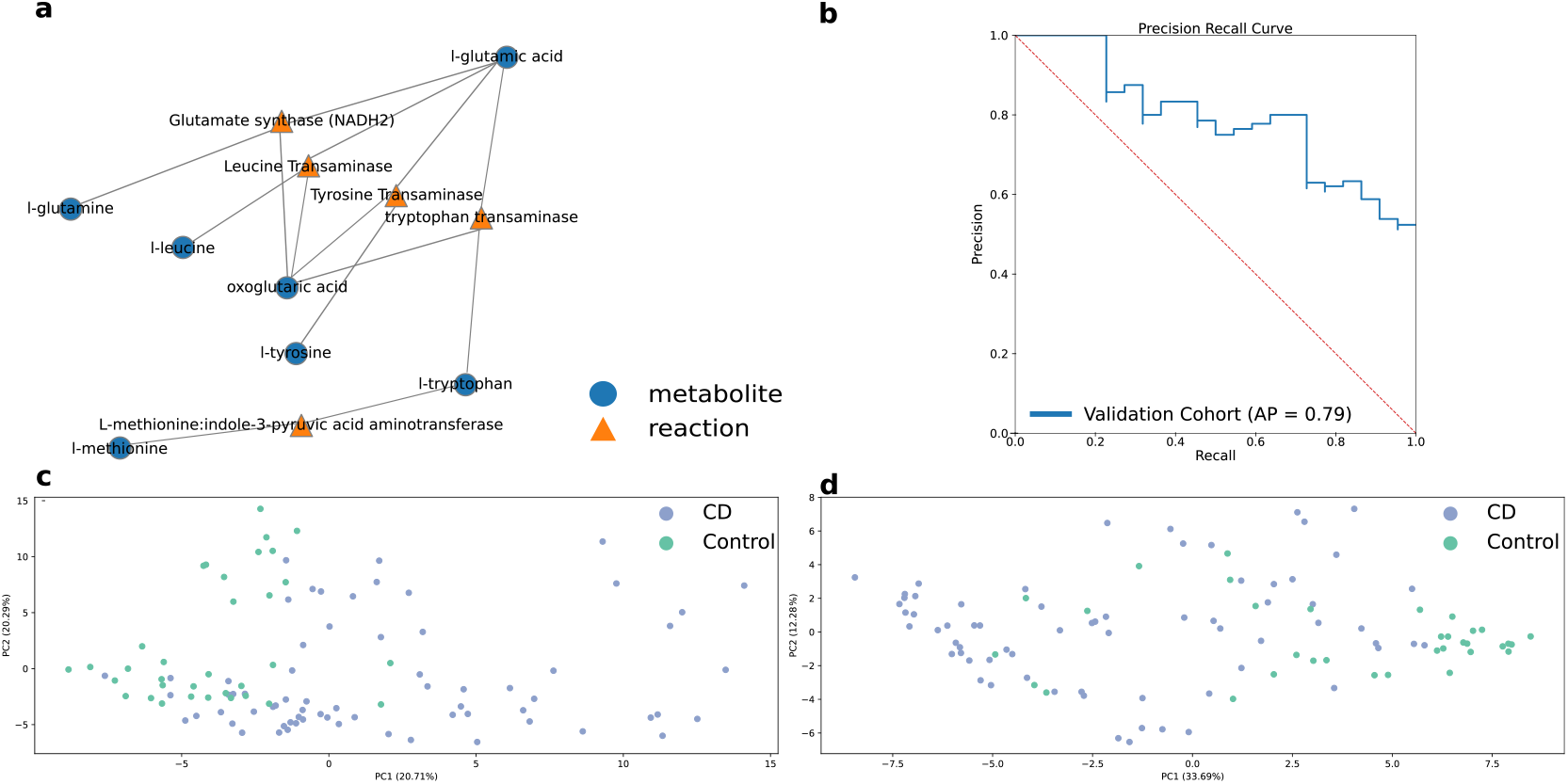
*mantra* results on Control vs. Crohn’s Disease stool samples from Franzosa et al. [30] **a** Subgraphs identified by the local search-based enrichment, repeated 5 times with different random starts. Two disconnected subgraphs are identified, one representing a part of cholic acid metabolism and the other one a part of amino acid metabolism. **b** Precision Recall (PR)-curve showing the predictive performance of a random forest model on the validation cohort trained on the PRISM cohort (both from Franzosa et al. [30]). The Expected performance by a random model is depicted by the red dashed line, the blue curve indicates the performance of the reaction activity data. With a PR-Area Under the Curve (AUC) of 0.79 the reaction estimate-based prediction seems to generalize well to the validation cohort. **c** PCA of the processed metabolite data. Samples colored by condition, which are mainly separated by PC1, explaining 20% of the variance in the dataset. Generally, control samples are more similar to each other than CD samples. **d** PCA on the basis of the reaction activity values computed with *mantra* with samples colored by condition. Even though PC1 explains a larger proportion of variance, the separation between sample groups is less clear than for the metabolome data.

To find a metabolically connected subgraph with high changes in reaction estimates between control and CD samples, we employed Simulated Annealing (SA) together with a local search, where the objective is to maximize the difference in distributions between control and CD samples (for details see Materials & Methods). This strategy is essentially an extension of the analysis presented in the previous section. Figure 3a, depicting the subgraph identified by the enrichment analysis, shows that exclusively amino acid interconversion reactions, mostly involving glutamate, are identified. Since Franzosa et al. [30] also performed metagenomics to quantify the differences in enzymatic capabilities in the gut lumen of control and CD samples, we checked whether any enzymes catalyzing the reactions in the subgraph are found to be significantly different (Supplementary Dataset 7 in [30]). Indeed, for 4 out of the 5 reactions, at least one enzyme capable of driving them is found to be differentially abundant between conditions (Supplementary Table S3). The only non-significant reaction, matched to Branched-chain-amino-acid transaminase, has a corrected q-value of 0.054, making it just barely missing the q-value cutoff of 0.05 used by the authors of [30]. With the general discussion about the choice of p-values and cutoffs in mind, one can conclude that the metabolic reactions identified with *mantra* are well reflected by the results of the metagenomics analysis.

The identified conversion between glutamine and glutamate is especially well-connected to IBD literature due to the role of glutamine-based signaling in the regulation of tissue integrity, inflammatory processes, and apoptosis [31]. The regulation of apoptosis is directly influenced by the conversion of glutamine to glutamate, which is then used to produce the reduced form of glutathione (GSH) together with cysteine and glycine [32, 33]. GSH is then used to regulate the redox potential via the binding of 2 GSH, resulting in the oxidized form of glutathione (GSSG). While only glutamate synthase was previously directly associated with IBD, tryptophan, tyrosine, and methionine metabolism have also been linked to CD [34, 35]. The results of this analysis in combination with the fact that all reactions are represented by differentially abundant enzymes, suggest that the subgraph identified by the local search algorithm on the basis of our proposed reaction activity metric yields a hypothesis consistent with additional measurements on expected enzyme levels and existing literature.

In addition to the PRISM cohort, which was used in the analyses presented above, Franzosa et al. [30] also introduce a validation cohort from a different hospital containing 22 control and 20 CD samples. We used this cohort to evaluate how well a predictive model trained on our reaction activity metric can generalize to a different cohort. The PR curve of this evaluation is shown in Figure 3b. The AUC of 0.79 demonstrates that the reaction activity estimation introduced is also generalizing across independent cohorts, making it suitable for clinical analyses. Additionally, the Receiver Operating Characteristic (ROC) curve for the same evaluation (AUC of 0.74) is shown in Supplementary Figure S3.

#### Associating Metabolic Reaction Activity with Microbial Species Abundances

A major advantage of using the residual variance instead of e.g. correlation coefficient is that they give per-sample estimates. Hence, they can be used to compute associations of metabolic reaction activity with features from other modalities. Computing all pairwise Spearman’s correlation coefficients between reaction values and microbial species resulted in 249 significant associations (q-value < 0.05) in the control group and 300 in the CD group, with 25 shared associations between the two groups. While these numbers appear rather low, the proportion of significant metabolite-microbe associations found by Franzosa et al. [30] is also below 10%. The correlation matrices and networks over all features are shown in Supplementary Figure S4. Since interpreting these networks without further analyses is a tedious task given their size and complex structure, we continue our analysis of multi-omics associations using the amino acid-reaction subgraph identified in the previous subsection (Figure 3a). More specifically, we look at the differences in correlations between control and CD patients. The rationale behind taking this approach is that a change in correlation potentially indicates a difference in the association between the features under the two conditions. In total, 35 microbial species have at least one significant association with a metabolic reaction that is different between the sample groups (Figure 4a). A large proportion of associations that are more positive in the control group are between a set of species and the three transaminase reactions, as well as the glutamate synthase reaction. The species with significant correlation changes to the methionine-related reaction, in contrast, are almost exclusively correlated with this reaction. Positively valued species are dominated by *Clostridium*, whereas those with negative values are dominated by *Streptococcus*.

**Figure 4:**
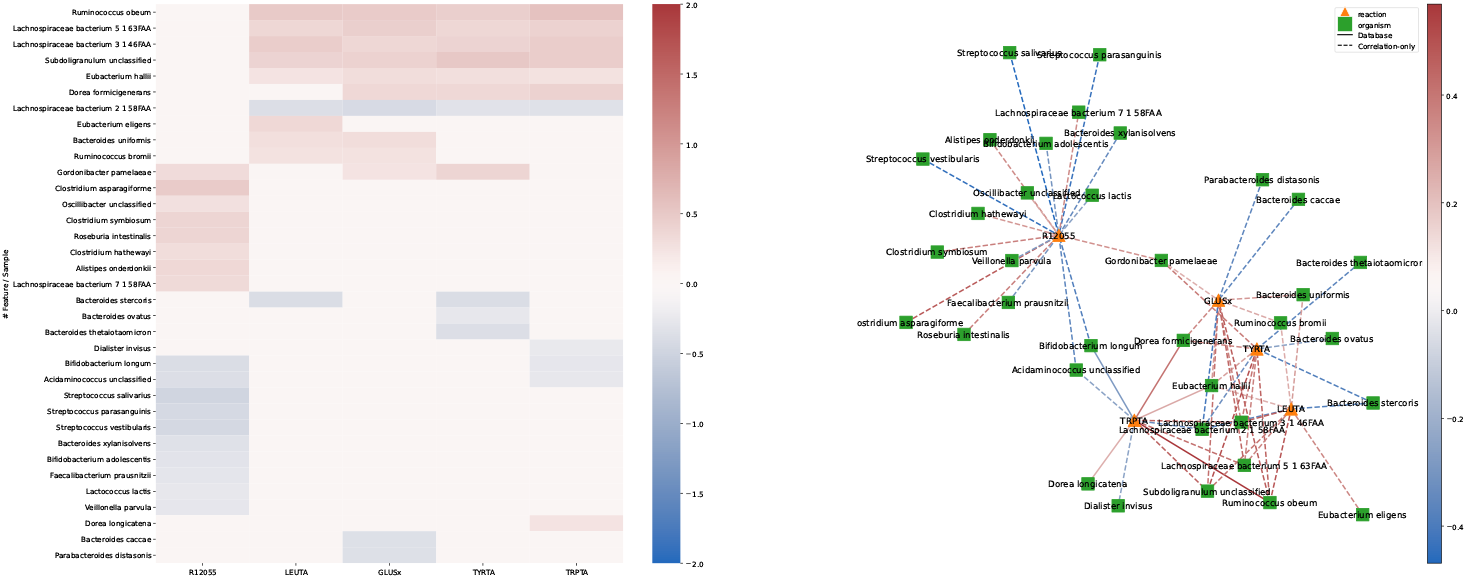
Spearman’s rank correlation-based associations between microbial species and enriched metabolic reactions. Reaction IDs matching the following reactions in Figure 3a: GLUSx, GLUSy, GLFRDOi: Glutamate synthase; LEUTA, LEUTAm: Leucine Transaminase; TYRTA, TYRTAm: Tyrosine Transaminase; TRPTA: tryptophan transaminase; R12055: methionine:indole-3-pyruvic acid aminotransferase. **a** Correlation matrix showing the difference in correlation coefficient between control and Crohn’s Disease (CD) samples for the reactions identified in the network enrichment and all microbial species. All correlation coefficients with a p-value > 0.05 are set to zero prior to computing the differences. **b** Correlation network resulting from the correlation matrix in a. Node color and shape indicated node type, edge color indicates the respective difference in correlation coefficient.

## 3 Discussion

In this work, we introduced an approach to estimate the changes in reaction activity between biological conditions using untargeted metabolomics data and metabolic networks. We demonstrate the ability of the proposed heuristic to recover known key reactions in biopsies of healthy and TNBC tissue and in stool samples of CD and non-IBD patients as a proof of concept. Furthermore, we showcase the possibility of computing multi-omics associations to these relative reaction activities via correlation metrics.

Despite the increasing use of metabolomics to study biological and biomedical phenomena, computational methods for a mechanistic interpretation of metabolomics data are rare. Especially metabolic reactions are only targeted when using metabolic modeling or when doing enrichment analyses using metabolite p-values (e.g. in [20]). The former is exact but often not feasible due to the requirements for data and model parameterization [36]. The latter option considers the biochemical connections between metabolites but disregards their quantitative relations and discards valuable information by discretizing with a threshold value. Inferring metabolic reaction activity without absolute quantification is hard, as values (i.e. MS intensities) are typically on metabolite-specific scales. Hence, even with a complete metabolic model, exact modeling is not possible. The approach we introduce circumvents these shortcomings by investigating the relative change of metabolite relations between sample groups on the basis of linear models. Based on the rationale that metabolites participating in the same active metabolic reaction show correlating properties [37], captured by the linear model, we propose that a change in the coefficients, describing the substrate-product relations, can be used as an indicator for a change in reaction activity. Following this idea, we use the residuals of two conditions for the same model as a proxy for this change of coefficients. The general applicability of our approach to generate mechanistic hypotheses is demonstrated by the results depicted in Figure 2d and Figure 3a and b, where we show that it pinpoints reactions by only using untargeted metabolomics data that were previously validated as key players in more complex experiments. Nevertheless, our approach may be limited with respect to non-linear behavior in reactions [37].

An advantage of using linear models is that correction of covariates, especially relevant in clinical studies, as well as the usage of regularized models such as elastic net [38] is simple and already implemented in the published code (see Code Availability Statement). Since each reaction is described by a separate model, our method considers each metabolic reaction in isolation, whereas in reality, reactions are connected through molecules participating in both reactions and many changes are propagated through the network. In cases where a consecutive sequence of reactions is actually changing, this might lead to the method only picking up the flanking reactions of this sequence. Additionally, in studies where the effect of external metabolite administration is “directly” given to the tissue/body site of sampling, like dietary interventions paired with gut/stool metabolomics, our method can be biased toward reactions in which the administered metabolites participate. Despite many disease-unrelated metabolic changes constantly happening, it is unlikely that such changes are picked up by the model. Especially in clinical studies, where high inter-sample heterogeneity is common, these effects will mostly remain as noise, whereas the true underlying changes are more constant across samples.

While this manuscript only evaluates application cases in a supervised form, the proposed idea can also be used in an unsupervised context. Therefore, applications such as *de-novo* subtyping need to be evaluated with respect to the robustness of similar metrics in future work. While the main focus of this work is the evaluation of a metabolic reaction activity metric, we also introduce a local search-based enrichment method to perform subgraph enrichment. Despite the good results, this approach is influenced by hyperparameter selection, such as temperature and allowed solution sizes for SA, and can become slow depending on the size of the network and the parameter settings. For example, the initial temperature *T*_0_ in SA controls the probability of accepting a random solution and thus the degree of exploration across the objective function landscape. Consequently, the choice of initial temperature is a trade-off between exploration and exploitation, and inappropriate temperature settings can lead to unstable or considerably sub-optimal solutions.

The participation of some metabolites in a large number of metabolic reactions may also be problematic from a graph-topological point of view due to their high connectivity within the metabolic network. Since our analysis does not include the neighborhood of a reaction, this does not affect the metric but only possible downstream applications acting on the network. However, it is not possible to distinguish reactions with the same substrates and products, as these will result in the same linear model, as well as the directionality of the reaction. In practical applications, this can mean that the hypothesis can include a larger number of possible enzyme candidates and, thus, more laborious hypothesis validation, even when the size of the results themselves is rather small.

Despite these limitations, the demonstrated ability of our method to accurately propose mechanistic hypotheses makes it a promising approach to improve the functional interpretation of metabolomics data in many experimental setups. By providing a metric for the quantitative approximation of reaction activity changes, it paves the way for a novel class of metabolomics data analysis methods. Given the wide-spread associations between functional metabolic changes in diseases [39, 40, 41, 42], we believe these developments can directly impact clinical research. In addition, the presented results give rise to the development of new strategies for prior knowledge-guided functional multi-omics integration to further strengthen biological and biomedical research on the level of metabolism.

## 4 Materials & Methods

### 4.1 Network Generation & ID mapping

To have a comprehensive database for human and microbial metabolism, we use a custom database. It contains the merged information from the Virtual Metabolic Human project [22], the KEGG [13], and the Reactome [21] database. The database is available through the provided Python package either through a public API or locally via a docker application provided (Code Availability Statement).

For a given metabolomics dataset, the identified metabolites are first mapped onto the internal database, either by directly using database identifiers, if given or by using the Metaboanalyst [43] name conversion API (http://api.xialab.ca/mapcompounds). Subsequently, the metabolic subnetwork containing all measured and mapped metabolites and all metabolic reactions for which at least one substrate and one product are measured is extracted. Metabolites not connected to any reaction in the subnetwork are removed.

### 4.2 Estimating Changes in Reaction Activity

The estimation of relative activity is based on reaction-wise linear models. This is based on findings from Krumsiek et al. [37] who showed that metabolites involved in active metabolic reactions have correlating properties. Prior to computing these models, all metabolites are mean-centered and scaled to unit variance to avoid scale-based biases in the residuals for each reaction. The input data is assumed to be normalized and transformed to follow (approximately) a normal distribution. Intuitively, our model tries to capture the change in relations between the substrates and products of a metabolic reaction between two conditions. The underlying assumption is that if reaction activity stays the same, so should the model coefficients, while they are expected to change if activity changes. Hence, while there can be similar correlation strength in both sample groups, the quantitative relation describing this correlation can change.

For each reaction in the metabolic network, the substrate abundances (*S*) are the predicting (i.e. independent) variables, and the product abundances (*P*) are the dependent variables. Therefore, each reaction model can be a multiple and/or multivariate regression, depending on the number of substrates and products that were measured for a particular reaction. The model for each reaction is formally defined as

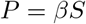

where *β* are the coefficients of the model and *P* and *S* are the intensities of the products and substrates of the respective reaction. Reactions with the same substrates and products, such as transport reactions, are skipped because *P* = *S*. The fitting of the model coefficients is happening in one of the sample groups. To avoid corruption of models by outliers, for each reaction, Cook’s distance [44] is computed for every sample. It is defined as the change of prediction relative to the model’s error when a sample is left out. For a sample *i* its Cook’s distance *D*_*i*_ is computed as

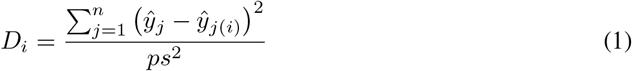

with ŷ _*j*_ and ŷ _*j*(*i*)_ being the predicted product intensities of the total model and the model without sample *i* respectively, for each product *j* of a reaction, *s*^2^ representing the model’s mean square error, and *p* the rank of the model. Generally, a high Cooks’s distance indicates a high influence of a sample and thus makes it more likely to be an outlier. If any sample has a distance value above a user-defined threshold, the model is fitted again without those samples. Alternatively, if no distance threshold is defined, we use the survival function of an F-distribution to provide p-values for selecting outliers to remove. Subsequently, the coefficient of determination (*R*^2^) is used to filter out all models that fail to describe the relation between substrate and product values adequately. Under the assumption that some metabolic reactions may only be active in one of the sample groups, this might mean that reactions are removed, which potentially describes a major difference between conditions. Therefore, we provide the option to re-compute models that fail in one group on samples from another group.

The reaction-wise models computed with the procedure explained above are then used to compute the reaction values for all samples. More specifically, the reaction activity estimates are calculated as the (normed) proportion of explained variance. For a sample *i* and a reaction *r* this estimate is defined as

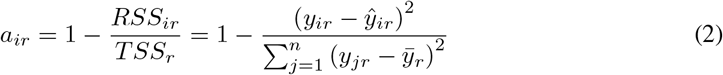

where ŷ represents the predicted product values and 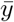 the mean product values. The summation in the denominator over *j* represents the total sum of squares (TSS), i.e. the residual sum of squares summed over all samples. Subsequently, the estimates are mean-centered and scaled to unit variance.

To obtain p-values describing how statistically different the activity estimate distributions between groups are, a Wilcoxon rank sum test is calculated for each reaction on the basis of the activity estimates computed as described before. The reported p-values are family-wise error rate (Bonferroni) corrected.

### 4.3 Multi-Omics Associations

To compute multi-omics associations, we use Spearman’s rank correlation implemented in the scipy package [45]. Corresponding p-values above a user-defined threshold are used to set the respective correlation coefficients to zero. Subsequently, all reactions and multi-omics features without a single significant correlation coefficient are removed. Correlations can either be computed for each group individually or over all samples.

### 4.4 Network Enrichment

The combinatorial optimization algorithm used for the network enrichment is an adapted version of a local search approach used in Rose and Köhler et al. [46]. Local search generally examines a search space in a greedy manner by iteratively testing local candidate solutions for the one with an optimal objective function. Candidate solutions are generated by applying one of three operations: node insertion, deletion, and substitution to the solution from the last iteration or a randomly selected subgraph in the first iteration.

While this procedure is sufficient to find local maxima, it cannot escape them, and thus, the resulting optimal solution is highly dependent on the initial starting point and the landscape of the objective function over the entire graph. Therefore, we use SA [47] to avoid stagnation and increase the chances of finding a global maximum. SA allows accepting non-optimal subgraphs with a probability decreasing with the number of iterations. This decrease is implemented through a temperature parameter *T* exponentially decaying by a rate *α* with the number of iterations *n*. Whenever the local search reaches a local maximum, a sub-optimal solution is accepted if

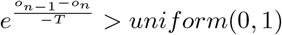

where *o*_*n*_ and *o*_*n*−1_ are the values of the objective function in the current and last iteration.

The objective function is defined as

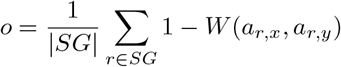

where *SG* indicates the set of vertices forming the current solution and *W* is the p-value of a Welch’s test comparing group *x* against group *y*. It is defined such that the local search maximizes the difference between the reaction activity estimate distributions between the two sample groups.

Even when using SA, finding a global optimum is not guaranteed and in some cases, multiple global optima might exist. Hence, the enriched subgraph is dependent on the randomly chosen initial solution. Therefore, our method runs the enrichment algorithm multiple times with different seeds and returns the union of all solutions. Another option instead of using the union would be the intersection of all repeats (an option in the package). In an intuitive sense, the union subgraph would give a lower probability of “false negatives” at the expense of a higher chance of including “false positives”. In practice, the choice also depends on the settings of other hyperparameters, such as the allowed solution size and the downstream experiments that the enrichment results should be used for.

### 4.5 Data Processing and Experiments

Common to both experiments presented in this manuscript is the usage of the networkx [48] and matplotlib [49] packages for handling network visualization and the scikit-learn library [50] for generating PCAs.

#### 4.5.1 Triple-Negative Breast Cancer Data Set

The metabolomics data provided by Xiao et al. [23] gives already normalized metabolite data. Therefore, only missing-value imputation with half of the feature-wise minimum, mean centering, and unit variance scaling was applied. To map the measured metabolites to our internal database, the HMDB [51] and KEGG [13] identifiers provided with the feature annotation were used together with a mapping database provided with the python package (see Code Availability Statement). Following the name mapping step, the data-specific metabolic network was also generated with package-provided functions using a neo4j database that we made publicly available (see Code Availability Statement).

For analyzing and visualizing the residual distributions, the scipy [45] and seaborn [52] libraries were used.

#### 4.5.2 Inflammatory Bowel Disease Data Set

For the metabolomics data from Franzosa et al. [30] missing values were imputed using half the feature-wise minimum. Subsequently, samples were quotient normalized [53] to account for sample-specific dilution effects and log-transformation was applied, such that features are approximately normally distributed. All features were then mean-centered and scaled to unit variance. To avoid leakage between the PRISM (discovery) and the validation cohort, the PRISM cohort was processed and parameters for each (parameter-dependent) step were retained. The validation cohort was then processed with the parameters from the discovery cohort.

Since the available data did not contain database identifiers, we used the Metaboanalyst [43] name conversion API (http://api.xialab.ca/mapcompounds) to obtain HMDB and KEGG IDs for all metabolites. The remaining name mapping and network generation steps are the same as described above for the other analyzed data set.

Microbial species data was imputed by setting all zero values to the total minimum divided by the number of zero values before applying centered log-ratio transformation [54].

For assessing the generalization of the predictive model using our reaction activity metric, the random forest implementation from the scikit-learn package [50] was used with the default parameter (no hyperparameter optimization was done). Reaction models were computed on the discovery, followed by training the classifier on the resulting values. Subsequently, PR and ROC curves were computed on the validation samples with reaction values estimated using the reaction models fitted on the discovery cohort control samples. The curves shown were also generated with scikit-learn and matplotlib [49] functions.

Multi-omics correlations were computed based on the computed reaction metrics and the processed microbial species data using Spearman’s correlation coefficient.

## Data Availability Statement

All data used in the experiments is publicly available from the referenced articles by Xiao et al. [23] and Franzosa et al. [30].

## Code Availability Statement

Source code including all presented experiments on GitHub https://github.com/lipitum/pymantra and Zenodo https://doi.org/10.5281/zenodo.13142372

Python package: https://pypi.org/project/pymantra

Database and API: https://github.com/lipitum/pymantraAPI

## Author Contributions

NK and JKP planned the work. NK designed and implemented the reaction activity estimation method and the network enrichment procedure. VW and NK parsed and merged the reaction databases. NK ran the evaluations. NK, JKP, VW and TDR wrote and reviewed the manuscript. JKP secured the funding. All authors read and accepted the manuscript in its final form.

## Acknowledgments

We thank Felix Niedermaier for helping with the database API setup and Emma Schonner for internal evaluations that are not part of this manuscript. This project was funded by the Bavarian State Ministry of Science and the Arts in the framework of the Bavarian Research Institute for Digital Transformation (bidt; JKP, VW, TDR, NK: Junior Research Group LipiTUM). TDR is supported through state funds approved by the State Parliament of Baden-Württemberg for the Innovation Campus Health + Life Science Alliance Heidelberg Mannheim.

## Conflicts of interest

The authors declare no conflicts of interest.

## Abbreviations

AUC: Area Under the Curve
CD: Crohn’s Disease
CMPK1: UMP-CMP kinase
CV: Cross Validation
ER: Estrogen Receptor
IBD: Inflammatory Bowel Disease
MS: Mass Spectrometry
PCA: Principal Component Analysis
PLS(-DA): Partial Least Squares(-Discriminant Analysis) PR Precision Recall
ROC: Receiver Operating Characteristic SA Simulated Annealing
TNBC: Triple-Negative Breast Cancer UC Ulcerative Colitits
UDP: Uridine Di-Phosphate
UGDH: UDP-glucose-dehydrogenase
UGT: UDP-glucuronosyltransferase
UMP: Uridine Mono-Phosphate

## Supplementary Figures

**Supplementary Figure S1:**
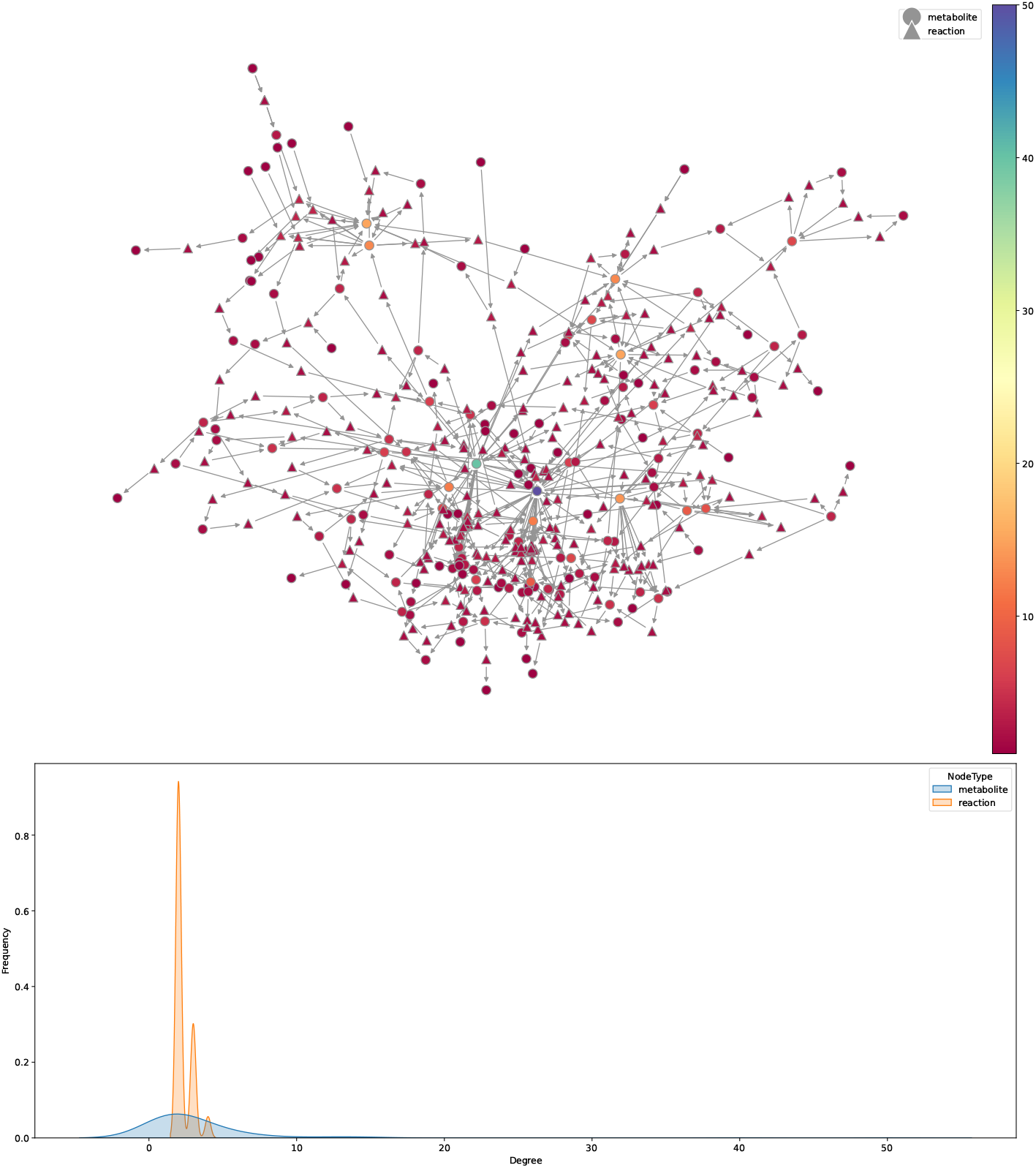
Node degrees of the metabolic network based on the data from Xiao et al. [23] **a** Metabolic network presented in Figure 2a showing the node degree by color. Node shapes indicate whether a node is a metabolite or a metabolic reaction. **b** Distributions of node degrees from a for metabolite and reaction nodes separately.

**Supplementary Figure S2:**
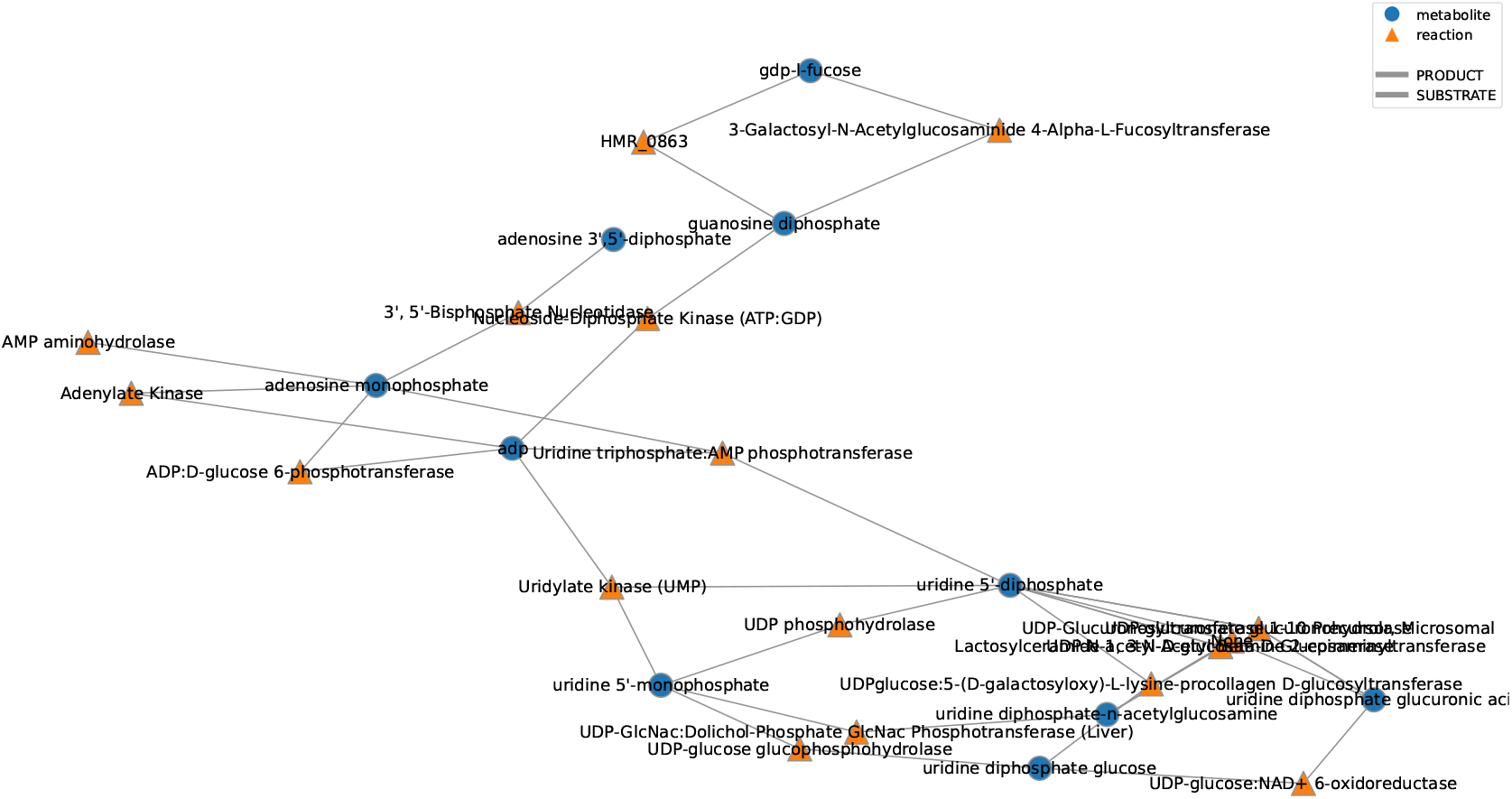
Induced subgraph using the reactions identified as significantly altered in activity in Figure 2d.

**Supplementary Figure S3:**
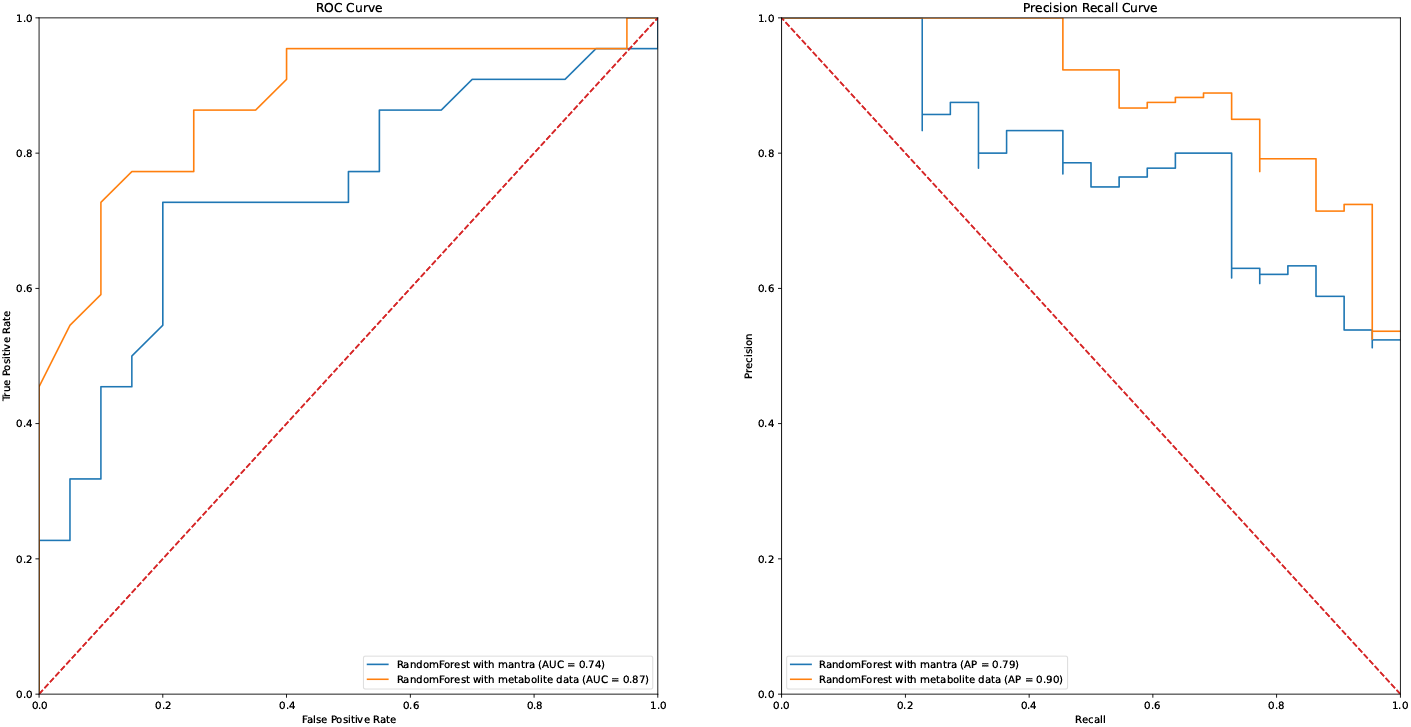
Receiver Operating Characteristic (ROC) and Precision Recall (PR) curves for the prediction of CD labels from the validation cohort of Franzosa et al. [30] on the processed metabolome data (orange) and the *mantra* estimates (blue).

**Supplementary Figure S4:**
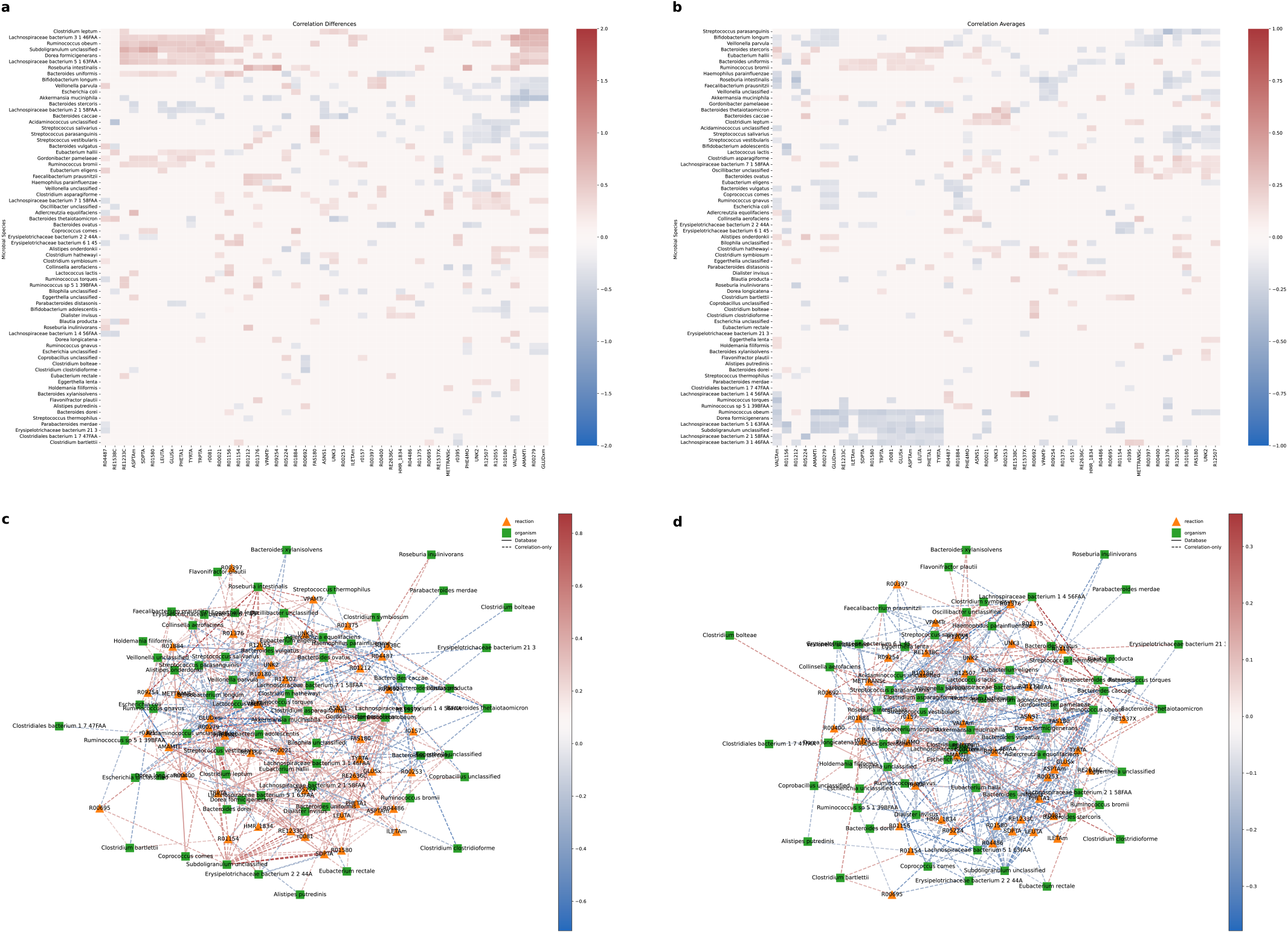
Significant correlations between metabolic reactions and microbial species in the PRISM cohort [30]. **a** Correlation matrix between metabolic reactions and microbial species showing the differences between non-IBD and CD. **b** Correlation matrix between metabolic reactions and microbial species showing the averages of non-IBD and CD. **c** Correlation network from the correlation differences in a. **d** Correlation network from the correlation averages in b.

## Supplemetary Tables

**Supplementary Table S1:**
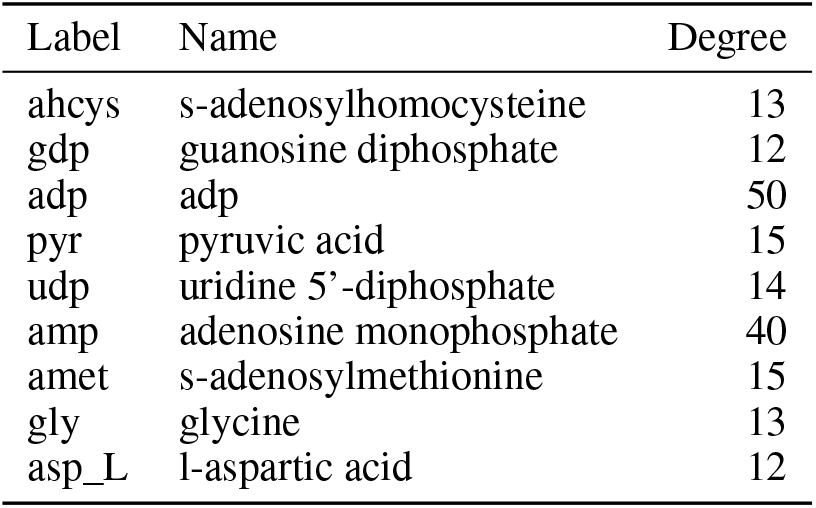
Node degrees of metabolites with a degree of at least 10 from the Triple-Negative Breast Cancer data set ([23])

| Label | Name | Degree |
| --- | --- | --- |
| ahcys | s-adenosylhomocysteine | 13 |
| gdp | guanosine diphosphate | 12 |
| adp | adp | 50 |
| pyr | pyruvic acid | 15 |
| udp | uridine 5'-diphosphate | 14 |
| amp | adenosine monophosphate | 40 |
| amet | s-adenosylmethionine | 15 |
| gly | glycine | 13 |
| asp_L | l-aspartic acid | 12 |

**Supplementary Table S2:** Detailed descriptions of the significant reactions from Figure 2d. The table is also available as a .csv file in the additional supplements.

| Short Name | Formula | Description |
| --- | --- | --- |
| URIDK1 | atp[c] + ump[c] -> adp[c] + udp[c] | Uridylate kinase (UMP) |
| BPNT | h2o[c] + pap[c] -> amp[c] + pi[c] | 3', 5'-Bisphosphate Nucleotidase |
| ADK1 | amp[c] + atp[c] <=> 2.0 adp[c] | Adenylate Kinase |
| UTPATPT | amp[c] + utp[c] <=> adp[c] + udp[c] | Uridine triphosphate:AMP phosphotransferase |
| R00155 | UDP + H2O <=> UMP + Orthophosphate | UDP phosphohydrolase |
| R00287 | UDP-glucose + H2O <=> UMP + D-Glucose 1-phosphate | UDP-glucose glucophosphohydrolase |
| NDPK1 | atp[c] + gdp[c] <=> adp[c] + gtp[c] | Nucleoside-Diphosphate Kinase (ATP:GDP) |
| GMPR | gmp[c] + 2.0 h[c] + nadph[c] -> imp[c] + nadp[c] | GMP Reductase |
| R00181 | AMP + H2O <=> IMP + Ammonia | AMP aminohydrolase |
| R08615 | UDP-glucuronate + H2O <=> UDP + D-Glucuronate | UDP-glucuronate glucuronohydrolase |
| R00414 | UDP-N-acetyl-alpha-D-glucosamine + H2O <=> N-Ac... | UDP-N-acetyl-D-glucosamine 2-epimerase |
| R00589 | L-Serine <=> D-Serine | serine racemase |
| HMR_0873 | fucgalfucgalcglgalgluside_hs[c] + gdp[c] + h[... | 3-Galactosyl-N-Acetylglucosaminide 4-Alpha-L-Fu... |
| R05327 | n UDP-N-acetyl-alpha-D-glucosamine + n UDP-gluc... | None |
| R00286 | UDP-glucose + H2O + 2 NAD+ <=> UDP-glucuronate ... | UDP-glucose:NAD+ 6-oxidoreductase |
| HMR_0863 | galacglcgalgluside_hs[c] + gdpfuc[c] -> gdp[c] ... | HMR_0863 |
| HMR_0859 | galgluside_hs[c] + uacgam[c] -> acglcgalgluside... | Lactosylceramide 1, 3-N-Acetyl-Beta-D-Glucosami... |
| R04491 | UDP-glucose + 5-(D-Galactosyloxy)-L-lysine-proc... | UDPglucose:5-(D-galactosyloxy)-L-lysine-procoll... |
| GLCNACPT_L | 0.1 dolp_L[c] + uacgam[c] -> 0.1 naglc2p_L[c] +... | UDP-GlcNac:Dolichol-Phosphate GlcNac Phosphotra... |
| UGT1A1r | estrone[r] + udpglcur[r] -> estroneglc[r] + udp... | UDP-Glucuronosyltransferase 1-10 Precursor, Mic... |
| R05804 | ADP + D-Glucose <=> AMP + D-Glucose 6-phosphate | ADP:D-glucose 6-phosphotransferase |

**Supplementary Table S3:** Comparing the reactions from the network enrichment results (Figure 2a) to metagenomics-based functional profiling. The table shows the name of the reaction displayed in the original subgraph plot together with the respective EC number and its respective corrected p-value (q-value) in Supplementary Dataset 7 in Franzosa et al. [30].

| Sub-graph reaction | Matched EC Number | q-value |
| --- | --- | --- |
| Glutamate syntase | 1.4.1.14 | 0.00497 |
| Tyrosine Transaminase | 2.6.1.57 | 0.00678 |
| tryptophan transaminase | 2.6.1.57 | 0.00678 |
| methionine:indole-3-pyruvic acid aminotransferase | 2.6.1.88 | 0.0126 |
| Leucine Transaminase | 2.6.1.42 | 0.0545 |

## Notes

### Competing Interest Statement

The authors have declared no competing interest.

### Summary of Updates

Corrected title and updated author information

## References

[1] David S. Wishart. “Metabolomics for Investigating Physiological and Pathophysiological Processes”. In: Physiological Reviews 99.4 (Oct. 2019). Publisher: American Physiological Society, pp. 1819–1875. DOI: 10.1152/physrev.00035.2018.

[2] Yue Xiong and Kun-Liang Guan. “Mechanistic insights into the regulation of metabolic enzymes by acetylation”. In: The Journal of Cell Biology 198.2 (July 2012), pp. 155–164. DOI: 10.1083/jcb.201202056.

[3] Abhisha Sawant Dessai, Poonam Kalhotra, Aaron T. Novickis, and Subhamoy Dasgupta. “Regulation of tumor metabolism by post translational modifications on metabolic enzymes”. en. In: Cancer Gene Therapy (Aug. 2022). Publisher: Nature Publishing Group, pp. 1–11. DOI: 10.1038/s41417-022-00521-x.

[4] Vincent J Hilser. “An ensemble view of allostery”. In: Science 327.5966 (2010), pp. 653–654.

[5] Luca Gerosa and Uwe Sauer. “Regulation and control of metabolic fluxes in microbes”. In: Current opinion in biotechnology 22.4 (2011), pp. 566–575.

[6] Janet E Lindsley and Jared Rutter. “Whence cometh the allosterome?” In: Proceedings of the National Academy of Sciences 103.28 (2006), pp. 10533–10535.

[7] Jeremy K Nicholson. “Global systems biology, personalized medicine and molecular epidemiology”. In: Molecular Systems Biology 2.1 (2006), p. 52. DOI: 10.1038/msb4100095.

[8] Claire L Gavaghan, Elaine Holmes, Eva Lenz, Ian D Wilson, and Jeremy K Nicholson. “An NMR-based metabonomic approach to investigate the biochemical consequences of genetic strain differences: application to the C57BL10J and Alpk: ApfCD mouse”. In: FEBS letters 484.3 (2000), pp. 169–174.

[9] Yang Chen, En-Min Li, and Li-Yan Xu. “Guide to Metabolomics Analysis: A Bioinformatics Workflow”. en. In: Metabolites 12.4 (Apr. 2022). Number: 4 Publisher: Multidisciplinary Digital Publishing Institute, p. 357. DOI: 10.3390/metabo12040357.

[10] Jianguo Xia and David S. Wishart. “MSEA: a web-based tool to identify biologically meaningful patterns in quantitative metabolomic data”. In: Nucleic Acids Research 38 (May 2010), W71–W77. DOI: 10.1093/nar/gkq329.

[11] Atanas Kamburov, Rachel Cavill, Timothy MD Ebbels, Ralf Herwig, and Hector C Keun. “Integrated pathway-level analysis of transcriptomics and metabolomics data with IMPaLA”. In: Bioinformatics 27.20 (2011), pp. 2917–2918.

[12] Adam Amara, Clément Frainay, Fabien Jourdan, Thomas Naake, Steffen Neumann, Elva María Novoa-Del-Toro, Reza M Salek, Liesa Salzer, Sarah Scharfenberg, and Michael Witting. “Networks and graphs discovery in metabolomics data analysis and interpretation”. In: Frontiers in Molecular Biosciences (2022), p. 223.

[13] Minoru Kanehisa, Yoko Sato, Masayuki Kawashima, Miho Furumichi, and Mao Tanabe. “KEGG as a reference resource for gene and protein annotation”. In: Nucleic acids research 44.D1 (2016), pp. D457–D462.

[14] Peter D Karp, Richard Billington, Ron Caspi, Carol A Fulcher, Mario Latendresse, Anamika Kothari, Ingrid M Keseler, Markus Krummenacker, Peter E Midford, Quang Ong, et al. “The BioCyc collection of microbial genomes and metabolic pathways”. In: Briefings in bioinformatics 20.4 (2019), pp. 1085–1093.

[15] Svetlana Volkova, Marta RA Matos, Matthias Mattanovich, and Igor Marín de Mas. “Metabolic modelling as a framework for metabolomics data integration and analysis”. In: Metabolites 10.8 (2020), p. 303.

[16] Song-Min Schinn, Carly Morrison, Wei Wei, Lin Zhang, and Nathan E Lewis. “Systematic evaluation of parameters for genome-scale metabolic models of cultured mammalian cells”. In: Metabolic Engineering 66 (2021), pp. 21–30.

[17] Olga Lazareva, Jan Baumbach, Markus List, and David B Blumenthal. “On the limits of active module identification”. In: Briefings in Bioinformatics 22.5 (2021), bbab066.

[18] Clément Frainay, Sandrine Aros, Maxime Chazalviel, Thomas Garcia, Florence Vinson, Nicolas Weiss, Benoit Colsch, Frédéric Sedel, Dominique Thabut, Christophe Junot, et al. “MetaboRank: network-based recommendation system to interpret and enrich metabolomics results”. In: Bioinformatics 35.2 (2019), pp. 274–283.

[19] Taher H Haveliwala. “Topic-sensitive pagerank”. In: Proceedings of the 11th international conference on World Wide Web. 2002, pp. 517–526.

[20] Sergio Picart-Armada, Francesc Fernández-Albert, Maria Vinaixa, Miguel A Rodríguez, Suvi Aivio, Travis H Stracker, Oscar Yanes, and Alexandre Perera-Lluna. “Null diffusion-based enrichment for metabolomics data”. In: PloS one 12.12 (2017), e0189012.

[21] Marc Gillespie, Bijay Jassal, Ralf Stephan, Marija Milacic, Karen Rothfels, Andrea Senff-Ribeiro, Johannes Griss, Cristoffer Sevilla, Lisa Matthews, Chuqiao Gong, et al. “The reactome pathway knowledgebase 2022”. In: Nucleic acids research 50.D1 (2022), pp. D687–D692.

[22] Alberto Noronha, Jennifer Modamio, Yohan Jarosz, Elisabeth Guerard, Nicolas Sompairac, German Preciat, Anna Dröfn Daníelsdóttir, Max Krecke, Diane Merten, Hulda S Haraldsdóttir, et al. “The Virtual Metabolic Human database: integrating human and gut microbiome metabolism with nutrition and disease”. In: Nucleic acids research 47.D1 (2019), pp. D614– D624.

[23] Yi Xiao, Ding Ma, Yun-Song Yang, Fan Yang, Jia-Han Ding, Yue Gong, Lin Jiang, Li-Ping Ge, Song-Yang Wu, Qiang Yu, et al. “Comprehensive metabolomics expands precision medicine for triple-negative breast cancer”. In: Cell Research 32.5 (2022), pp. 477–490.

[24] Rachel Sparks, Cornelia M. Ulrich, Jeannette Bigler, Shelley S. Tworoger, Yutaka Yasui, Kumar B. Rajan, Peggy Porter, Frank Z. Stanczyk, Rachel Ballard-Barbash, Xiaopu Yuan, Ming Gang Lin, Lynda McVarish, Erin J. Aiello, and Anne McTiernan. “UDP-glucuronosyltransferase and sulfotransferase polymorphisms, sex hormone concentrations, and tumor receptor status in breast cancer patients”. In: Breast Cancer Research 6.5 (June 2004), R488. DOI: 10.1186/bcr818.

[25] Bao-Xia He, Bin Qiao, Alfred King-Yin Lam, Xiu-Li Zhao, Wen-Zhou Zhang, and Hui Liu. “Association between UDP-glucuronosyltransferase 2B7 tagSNPs and breast cancer risk in Chinese females”. en. In: Clinical and Experimental Pharmacology and Physiology 45.5 (2018), pp. 437–443. DOI: 10.1111/1440-1681.12908.

[26] Ning Qing Liu, Tommaso De Marchi, Annemieke Timmermans, Anita M. A. C. Trapman-Jansen, Renée Foekens, Maxime P. Look, Marcel Smid, Carolien H. M. van Deurzen, Paul N. Span, Fred C. G. J. Sweep, Julie Benedicte Brask, Vera Timmermans-Wielenga, John A. Foekens, John W. M. Martens, and Arzu Umar. “Prognostic significance of nuclear expression of UMP-CMP kinase in triple negative breast cancer patients”. en. In: Scientific Reports 6.1 (Aug. 2016). Number: 1 Publisher: Nature Publishing Group, p. 32027. DOI: 10.1038/srep32027.

[27] Daiana L. Vitale, Ilaria Caon, Arianna Parnigoni, Ina Sevic, Fiorella M. Spinelli, Antonella Icardi, Alberto Passi, Davide Vigetti, and Laura Alaniz. “Initial Identification of UDP-Glucose Dehydrogenase as a Prognostic Marker in Breast Cancer Patients, Which Facilitates Epirubicin Resistance and Regulates Hyaluronan Synthesis in MDA-MB-231 Cells”. en. In: Biomolecules 11.2 (Feb. 2021). Number: 2 Publisher: Multidisciplinary Digital Publishing Institute, p. 246. DOI: 10.3390/biom11020246.

[28] Satu Tiainen, Sanna Oikari, Markku Tammi, Kirsi Rilla, Kirsi Hämäläinen, Raija Tammi, Veli-Matti Kosma, and Päivi Auvinen. “High extent of O-GlcNAcylation in breast cancer cells correlates with the levels of HAS enzymes, accumulation of hyaluronan, and poor outcome”. en. In: Breast Cancer Research and Treatment 160.2 (Nov. 2016), pp. 237–247. DOI: 10.1007/s10549-016-3996-4.

[29] Xiaochuan Liu, Gwendolyn Gonzalez, Xiaoxia Dai, Weili Miao, Jun Yuan, Ming Huang, David Bade, Lin Li, Yuxiang Sun, and Yinsheng Wang. “Adenylate Kinase 4 Modulates the Resistance of Breast Cancer Cells to Tamoxifen through an m6A-Based Epitranscriptomic Mechanism”. English. In: Molecular Therapy 28.12 (Dec. 2020). Publisher: Elsevier, pp. 2593– 2604. DOI: 10.1016/j.ymthe.2020.09.007.

[30] Eric A Franzosa, Alexandra Sirota-Madi, Julian Avila-Pacheco, Nadine Fornelos, Henry J Haiser, Stefan Reinker, Tommi Vatanen, A Brantley Hall, Himel Mallick, Lauren J McIver, et al. “Gut microbiome structure and metabolic activity in inflammatory bowel disease”. In: Nature microbiology 4.2 (2019), pp. 293–305.

[31] Min-Hyun Kim and Hyeyoung Kim. “The Roles of Glutamine in the Intestine and Its Implication in Intestinal Diseases”. en. In: International Journal of Molecular Sciences 18.5 (May 2017). Number: 5 Publisher: Multidisciplinary Digital Publishing Institute, p. 1051. DOI: 10.3390/ijms18051051.

[32] Erich Roth, Rudolf Oehler, Nicole Manhart, Ruth Exner, Barbara Wessner, Eva Strasser, and Andreas Spittler. “Regulative potential of glutamine—relation to glutathione metabolism”. In: Nutrition 18.3 (2002), pp. 217–221.

[33] AG Hall. “The role of glutathione in the regulation of apoptosis.” In: European journal of clinical investigation 29.3 (1999), pp. 238–245.

[34] Almut Heinken, Johannes Hertel, and Ines Thiele. “Metabolic modelling reveals broad changes in gut microbial metabolism in inflammatory bowel disease patients with dysbiosis”. en. In: npj Systems Biology and Applications 7.1 (May 2021). Number: 1 Publisher: Nature Publishing Group, pp. 1–11. DOI: 10.1038/s41540-021-00178-6.

[35] Xochitl C. Morgan, Timothy L. Tickle, Harry Sokol, Dirk Gevers, Kathryn L. Devaney, Doyle V. Ward, Joshua A. Reyes, Samir A. Shah, Neal LeLeiko, Scott B. Snapper, Athos Bousvaros, Joshua Korzenik, Bruce E. Sands, Ramnik J. Xavier, and Curtis Huttenhower. “Dysfunction of the intestinal microbiome in inflammatory bowel disease and treatment”. In: Genome Biology 13.9 (Sept. 2012), R79. DOI: 10.1186/gb-2012-13-9-r79.

[36] Amit Rai and Kazuki Saito. “Omics data input for metabolic modeling”. In: Current opinion in biotechnology 37 (2016), pp. 127–134.

[37] Jan Krumsiek, Karsten Suhre, Thomas Illig, Jerzy Adamski, and Fabian J Theis. “Gaussian graphical modeling reconstructs pathway reactions from high-throughput metabolomics data”. In: BMC systems biology 5 (2011), pp. 1–16.

[38] Hui Zou and Trevor Hastie. “Regularization and variable selection via the elastic net”. In: Journal of the royal statistical society: series B (statistical methodology) 67.2 (2005), pp. 301– 320.

[39] Khushboo G Upadhyay, Devendra C Desai, Tester F Ashavaid, and Alpa J Dherai. “Microbiome and metabolome in inflammatory bowel disease”. In: Journal of Gastroenterology and Hepatology 38.1 (2023), pp. 34–43.

[40] Daniel R Schmidt, Rutulkumar Patel, David G Kirsch, Caroline A Lewis, Matthew G Vander Heiden, and Jason W Locasale. “Metabolomics in cancer research and emerging applications in clinical oncology”. In: CA: a cancer journal for clinicians 71.4 (2021), pp. 333–358.

[41] Vivian Tounta, Yi Liu, Ashleigh Cheyne, and Gerald Larrouy-Maumus. “Metabolomics in infectious diseases and drug discovery”. In: Molecular Omics 17.3 (2021), pp. 376–393.

[42] Christopher B Newgard. “Metabolomics and metabolic diseases: where do we stand?” In: Cell metabolism 25.1 (2017), pp. 43–56.

[43] Jasmine Chong, David S Wishart, and Jianguo Xia. “Using MetaboAnalyst 4.0 for comprehensive and integrative metabolomics data analysis”. In: Current protocols in bioinformatics 68.1 (2019), e86.

[44] R Dennis Cook. “Detection of influential observation in linear regression”. In: Technometrics 19.1 (1977), pp. 15–18.

[45] Pauli Virtanen, Ralf Gommers, Travis E. Oliphant, Matt Haberland, Tyler Reddy, David Cournapeau, Evgeni Burovski, Pearu Peterson, Warren Weckesser, Jonathan Bright, Stéfan J. van der Walt, Matthew Brett, Joshua Wilson, K. Jarrod Millman, Nikolay Mayorov, Andrew R. J. Nelson, Eric Jones, Robert Kern, Eric Larson,J J Carey, İlhan Polat, Yu Feng, Eric W. Moore, Jake VanderPlas, Denis Laxalde, Josef Perktold, Robert Cimrman, Ian Henriksen, E. A. Quintero, Charles R. Harris, Anne M. Archibald, Antônio H. Ribeiro, Fabian Pedregosa, Paul van Mulbregt, and SciPy 1.0 Contributors. “SciPy 1.0: Fundamental Algorithms for Scientific Computing in Python”. In: Nature Methods 17 (2020), pp. 261–272. DOI: 10.1038/s41592-019-0686-2.

[46] Tim D Rose, Nikolai Köhler, Lisa Falk, Lucie Klischat, Olga E Lazareva, and Josch K Pauling. “Lipid network and moiety analysis for revealing enzymatic dysregulation and mechanistic alterations from lipidomics data”. In: Briefings in Bioinformatics 24.1 (2023), bbac572.

[47] Peter JM Van Laarhoven, Emile HL Aarts, Peter JM van Laarhoven, and Emile HL Aarts. Simulated annealing. Springer, 1987.

[48] Aric Hagberg, Pieter Swart, and Daniel S Chult. Exploring network structure, dynamics, and function using NetworkX. Tech. rep. Los Alamos National Lab.(LANL), Los Alamos, NM (United States), 2008.

[49] J. D. Hunter. “Matplotlib: A 2D graphics environment”. In: Computing in Science & Engineering 9.3 (2007), pp. 90–95. DOI: 10.1109/MCSE.2007.55.

[50] F. Pedregosa, G. Varoquaux, A. Gramfort, V. Michel, B. Thirion, O. Grisel, M. Blondel, P. Prettenhofer, R. Weiss, V. Dubourg, J. Vanderplas, A. Passos, D. Cournapeau, M. Brucher, M. Perrot, and E. Duchesnay. “Scikit-learn: Machine Learning in Python”. In: Journal of Machine Learning Research 12 (2011), pp. 2825–2830.

[51] David S Wishart, AnChi Guo, Eponine Oler, Fei Wang, Afia Anjum, Harrison Peters, Raynard Dizon, Zinat Sayeeda, Siyang Tian, Brian L Lee, et al. “HMDB 5.0: the human metabolome database for 2022”. In: Nucleic Acids Research 50.D1 (2022), pp. D622–D631.

[52] Michael L. Waskom. “seaborn: statistical data visualization”. In: Journal of Open Source Software 6.60 (2021), p. 3021. DOI: 10.21105/joss.03021.

[53] Frank Dieterle, Alfred Ross, Götz Schlotterbeck, and Hans Senn. “Probabilistic quotient normalization as robust method to account for dilution of complex biological mixtures. Application in 1H NMR metabonomics”. In: Analytical chemistry 78.13 (2006), pp. 4281–4290.

[54] John Aitchison. “The statistical analysis of compositional data”. In: Journal of the Royal Statistical Society: Series B (Methodological) 44.2 (1982), pp. 139–160.

